# First whole-genome sequence of *Cervid atadenovirus A* outside of the United States from an Adenoviral hemorrhagic disease epizootic of black-tailed deer in Canada

**DOI:** 10.1101/2022.02.10.479879

**Authors:** Oliver Lung, Mathew Fisher, Michelle Nebroski, Glenna McGregor, Helen Schwantje, Tomy Joseph

## Abstract

A complete 30,616 nucleotide *Cervid atadenovirus A* genome was determined from tissues of a black-tailed deer that died in 2020 in British Columbia, Canada. Unique, non-synonymous SNPs in the E1B, Iva2 and E4.3 coding regions, and deletions totalling 74 nucleotides not observed in moose and red deer isolates were present.

## Main Text

Adenovirus hemorrhagic disease (AHD) is an acute, infectious, and usually fatal viral disease of several cervid species. AHD was first described in black-tailed deer in California in 1993; although, retrospective analysis of tissues suggests AHD was present in California in 1981 (1). The disease was previously reported in Canada (2), although no genome sequences are available. Deer mortality was reported on Galiano Island, British Columbia in September 2020. Analyses in BC and at UC Davis confirmed AHD. Since then, hundreds of black-tailed deer on many Gulf Islands and Vancouver Island have died of confirmed AHD. The disease was also observed in black-tailed deer in neighbouring Washington state in 2021.

There are two forms of AHD: a systemic form, primarily observed in the deer in BC, and a localized form (3). Systemic disease is characterized by fulminant pulmonary edema and intestinal hemorrhage, while the localized form is characterized by localized vasculitis, ulceration and necrosis in the mouth and stomachs (3). Adenoviruses are non-enveloped viruses with linear dsDNA genomes that have worldwide distribution and infect a wide range of vertebrates (4). Members of the family *Adenoviridae* are currently divided into six genera (4), including *Atadenovirus* with ten recognized species known to infect birds, ruminants, reptiles and marsupials (5, 6). Species demarcation within *Atadenovirus* is determined by attributes including phylogenetic distance (>10-15%), host range, cross-neutralization, and gene organization at the right end of the genome (6). The ten currently available complete genomes for *deer atadenovirus A* (Odocoileus adenovirus 1; OdAdV-1), the causative agent for AHD, were all obtained from cervids in the United States. These genomes were from moose, red deer, mule deer, black-tailed deer, white-tailed deer, and Rocky Mountain elk (Figure 1c). Renaming of OdAdV-1 to *cervid atadenovirus A* has been proposed (7), but not yet formally adopted.

**Figure 1.**
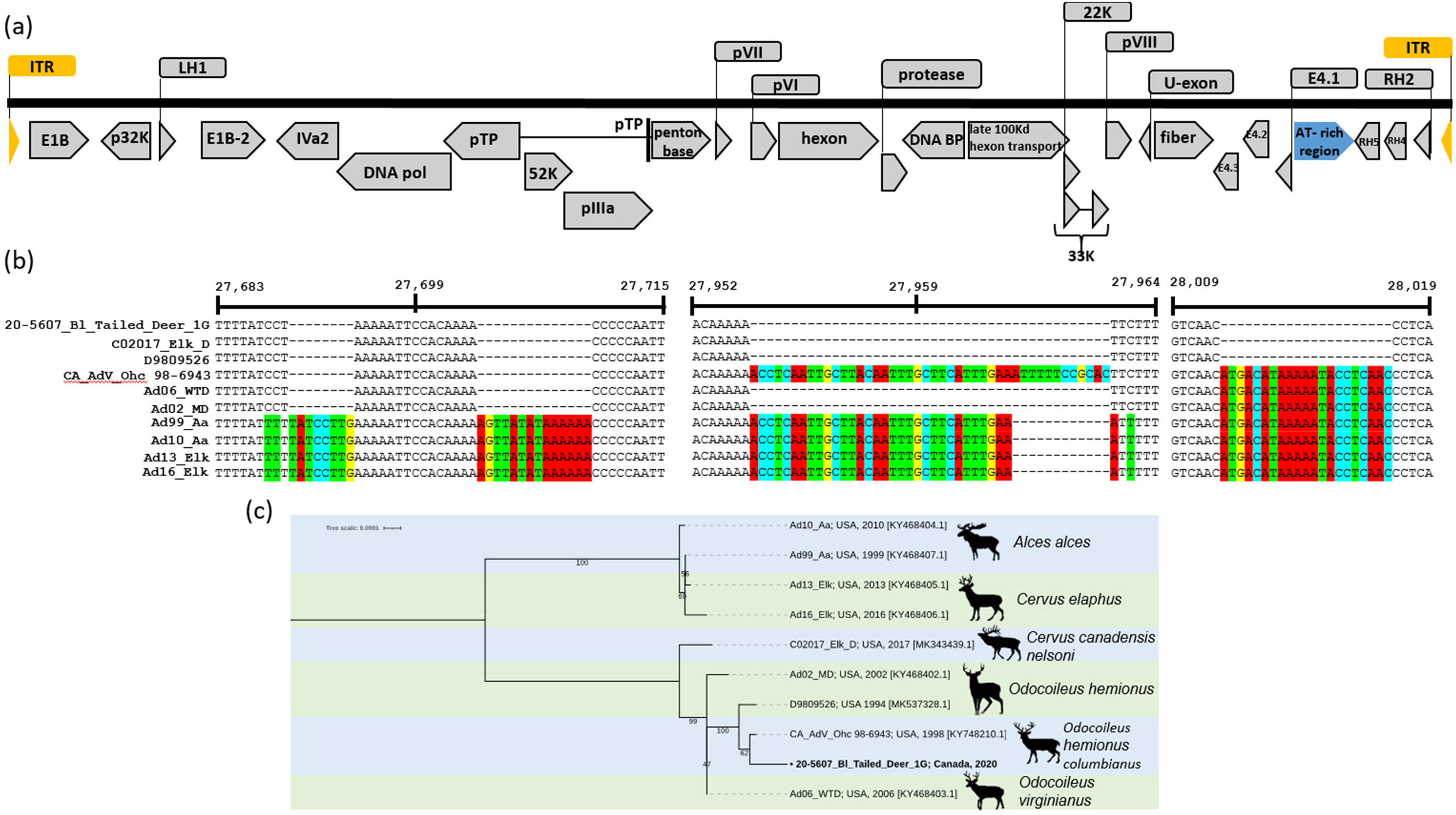
(a) Genome organization of *cervid atadenovirus A*. (b) Nucleotide alignment of the variable region of the AT-rich non-coding region. Insertions and SNPs relative to the 20-5607_Bl_Tailed_Deer_1G genome are highlighted, and genome positions are also relative to 20-5607_Bl_Tailed_Deer_1G. (c) Mid-point rooted maximum likelihood phylogenetic tree of *cervid atadenovirus A* (deer atadenovirus A) whole genomes. Alignment was performed using MAFFT (9), tree was generated with IQ-TREE (11) on find best model setting with ModelFinder (12) (model HKY-F chosen) and 1,000 ultra-fast bootstraps (13). Isolate names of sequences as well as location and year of collection date are shown and species name of host is indicated on the right. The genome presented here is indicated with a “•”.

High-throughput sequencing and analysis were performed as previously described (8). Contigs with homology to cervid atadenoviruses were observed in all three sequenced nasal swab samples. Adenovirus reads represented 13,855 of 6,536,978 (0.2%), 479,980 of 7,576,032 (6.3%) and 566,713 of 9,124,356 total reads (6.2%). Mean coverage depths were 106x, 3995x and 4521x, respectively. The resulting sequences, aligned using MAFFT (9), were identical in overlapping regions, and therefore combined into a single sequence. The presence of full inverted terminal repeats (ITR) indicated the genome was complete.

The assembled genome was 30,616 bp. Comparison with the reference (KY748210.1) shows 99.96% pairwise nucleotide identity, including 100% identity in the ITR. A SNP causing a potential 12 amino acid truncation in the viral replication-critical E1B protein (10) and non-synonymous SNPs in the Iva2 (T5486C) and E4.3 (C25817T) proteins were observed. Remaining SNPs were either synonymous or in non-coding regions. The variable region of the non-coding A/T-rich region contained four deletions totalling 74 nucleotides compared to isolates from moose and red deer, and appears to cluster with Rocky Mountain elk and mule deer isolates (Figure 1). Phylogenetic analysis shows our sequence is most closely related to the other black-tailed deer sequence available (KY748210.1), sampled in the Kentucky in 1998.

## Data availability

Nucleotide sequences including annotations and raw sequencing reads were deposited in GenBank and the Sequence Read Archive (SRA), National Center for Biotechnology Informations with the following GenBank and SRA accession numbers respectively: OM470968 and SAMN25644053. The version described in this paper is the first version.

## Acknowledgements

MN was supported in part by the LabsCanada Equipment Sharing and Scientific Platforms pilot project. The authors acknowledge Dr. Oksana Vernygora and Jossip Rudar for feedback on the manuscript.

## Conflict of Interest

The authors declare no conflict of interest.

## Author contribution

OL and TJ conceived the study. MF prepared the NGS sequencing library. MF, MN and OL analyzed the data. HS coordinated sample collection and submission and TJ and GM performed laboratory analyses. All co-authors contributed to the manuscript.

## Notes

### Competing Interest Statement

The authors have declared no competing interest.

